# Investigation of the Relationship Between Calpain and HMGB1/TLR4/NF-KB Signalling Pathway in Multiple Sclerosis and Other Demyelinating Diseases

**DOI:** 10.1101/2024.02.02.578379

**Authors:** Firdevs Uluc, Sule Aydin Turkoglu, Bihter Gokce Celik, Seyda Karabork, Seyit Ali Kayis

## Abstract

**Background:** To develop more effective treatments for demyelinating diseases, it is essential to identify the associated signaling pathways and factors. The objective of this study was to investigate the possible correlation between Calpain-1 (CAPN1) and Calpain-2 (CAPN2) with the HMGB1/TLR4/NF-κB signaling pathway and to evaluate the influence of these proteins on Interleukin 17A (IL-17A) and Interleukin 37 (IL-37) cytokines in individuals with newly diagnosed and untreated Multiple Sclerosis (MS) and Neuromyelitis Optica Spectrum Disorder (NMOSD).

**Methods:** In this pilot study, a total of 73 newly diagnosed patients were recruited, including 36 with MS, 9 with NMOSD, and an unhealthy control group composed of 28 individuals with Pseudotumour cerebri (PTC). To ensure accuracy and transparency, all groups’ demographic and clinical characteristics were meticulously described. ELISA technique was utilized to compare levels of CAPN1, CAPN2, HMGB1, TLR4, and NF-KB, as well as IL-17A and IL-37 cytokines, between the case and the unhealthy control groups. The expectation from these findings is to provide valuable insights into the pathophysiological mechanisms of these neurological disorders, possibly opening the door to novel therapeutic perspectives.

**Results:** In patients with MS, the levels of CAPN1 were found to be higher than those in patients with NMOSD and PTC. Similarly, the level of CAPN2 was significantly higher in patients with MS than in patients with NMOSD and higher in patients with PTC than in patients with NMOSD. There were no differences in the levels of HMGB1, TLR4, NF-κB, IL-17A, and IL-37 between the groups. Age and gender did not affect any of the parameters. In the MS group, both CAPN1 and CAPN2 showed positive correlation with HMGB1, TLR4, and NF-κB levels.

**Conclusions:** It may be suggested that CAPN1 may exhibit greater efficacy than CAPN2 during the initial stages of neuroinflammation. To obtain deeper and more guiding results of the varying levels of CAPN1 and CAPN2, and their relationship with the HMGB1/TLR4/NF-κB signaling pathway, it is advisable to conduct *in-vivo* and *in-vitro* prospective studies featuring CAPN1-specific inhibitors with larger study groups.

## 1. Introduction

MS is a chronic and inflammatory disease that affects the central nervous system (CNS). The degeneration and demyelination of neurons, which can lead to axonal loss and neuronal damage are the main characteristics of this disease (Yamout & Alroughani, 2018). This complex disease is known to cause a variety of symptoms that can affect the autonomic, motor, visual, and sensory systems. Over time, patients may experience episodes of acute neurological impairment or deterioration. MS is generally observed in young adults aged 20-40 years and affects approximately 2.3 million people worldwide. The prevalence ranges from 50-300 per 100000 people, with a higher incidence in women with a ratio of 1.5:1 to 2.5:1 (Thompson et al., 2018). The underlying causes of the disease remain not yet fully understood, however, the incidence and prevalence of MS are increasing worldwide day by day (Browne et al., 2014).

Neuromyelitis optica spectrum disorder (NMOSD) is a rare disease that primarily affects the spinal cord and optic nerve, as well as other parts of the brain. It was previously unclear whether NMOSD was a distinct disease or a more severe form of MS, but with the identification of the Aquapoin-4 (AQP4) water channel, NMOSD is now defined as a different disease from MS with its diagnostic criteria and treatments (Araki & Yamamura, 2017; Weinshenker & Wingerchuk, 2017).

Understanding the clinical manifestations and disease course of CNS demyelinating diseases requires knowledge of direct targets and signaling pathways, given their heterogeneous immunopathology (Popescu & Lucchinetti, 2012). To effectively treat these complex diseases, therapeutic agents must address both inflammation and neurodegeneration (W. Smith et al., 2017). The calpain system offers promise in this regard, with its ability to directly impact both aspects.

Calpains are a group of nonlysosomal cysteine proteases that are calcium-dependent and expressed in various cells of humans and other organisms. Their roles in various physiological and pathological processes are reported (Mahaman et al., 2019). The most extensively studied subtypes of the calpains are CAPN1 and CAPN2, which are also known as μ-calpain and m-calpain, respectively. These are widely expressed in most mammalian organs and cells; but are predominantly found in the brain (Sorimachi et al., 2012; Wang et al., 2018).

Calpains are naturally occurring enzymes found in the cytosol as a pro-enzyme. In resting cells, the normal range of free Ca^2+^ concentrations is typically between 50-100 nM. Within the normal range of intracellular free Ca^2+^ concentrations, calpain performs several important physiological functions including cytoskeletal reorganization, long-term potentiation, hormone processing, and protein turnover (Sorimachi et al., 2012; Wang et al., 2018). When intracellular free Ca^2+^ concentrations increase, the pro-enzyme is activated and can also translocate upon contact with membrane-bound phospholipids for the degradation of membrane proteins (W. Smith et al., 2017) and uncontrolled and prolonged calpain activation may occur (Baraban et al., 2018; Yamashima, 2016).

HMGB1 is a protein that exists in all mammalian cells and does not bind to histone DNA. Its primary structure consists of two homologous HMG box domains, A box, and B box, as well as a C-terminal acidic tail (Sternberg et al., 2016; Štros, 2010). This protein is present in astrocytes, neurons, and microglia in both the CNS and peripheral nervous system, as well as in the cytoplasm and extracellular spaces of neurons and Schwann cells (Fang et al., 2012). Furthermore, it has a considerable expression in regions of brain neurogenesis, such as the dentate gyrus and olfactory bulb (Guazzi et al., 2003). HMGB1 has a critical role as a DNA chaperone in the nucleus during normal physiological conditions. In macrophages, it is actively secreted or passively released from damaged or dead cells. Additionally, it has a pivotal function as a cytokine-inducing early inflammatory response. HMGB1 carries out its physiological role by binding to various membrane receptors, including receptors for advanced glycation end products (RAGE) and toll-like receptors (TLRs), which initiate the signaling cascade. When HMGB1 interacts with TLR4, it triggers the upregulation of NF-κB and results in the release of proinflammatory cytokines, chemokines, and enzymes (Li et al., 2018; Trotta et al., 2014). Notably, the release of pro-inflammatory cytokines by macrophages in response to HMGB1 depends on the presence of TLR4 (Harris et al., 2012).

The molecular mechanisms that are responsible for inflammation and immune response in the development of MS and other demyelinating diseases are not yet fully understood. It is crucial to determine these mechanisms to develop effective treatment options for this disease. Although there are various treatment alternatives available for patients with MS and NMOSD in the active phase, their short-term and limited efficacy, side effects, and inability to prevent relapses make it necessary to develop advanced treatment options that are specific to each of these diseases. According to a recent study, patients with Systemic Sclerosis, an autoimmune disease, showed a correlation between calpain and HMGB1 levels (Zheng et al., 2020). This study builds on that idea, suggesting that CAPNs and HMGB1, also known as platelet-derived microparticles, may be related to MS and other demyelinating diseases. By examining the relationship between CAPN1 and CAPN2 with HMGB1 levels, as well as the differences in protein levels in these two pathologies, we hope to develop a unique perspective for understanding both inflammation and demyelination processes in MS and NMOSD.

## 2. Methods

### 2.1 Subjects

The study received ethical approval from the Bolu Abant Izzet Baysal University (BAIBU) Local Ethical Committee under decision number 2021/305 on April 1st, 2022, and followed the guidelines outlined in the Helsinki Declaration. Cerebrospinal fluid samples (CSF) from seventy-three patients who were diagnosed between February 2021 and May 2023 at the Neurology outpatient clinic of the BAIBU Training and Research Hospital in Bolu, Türkiye were used. The case group included thirty-six patients diagnosed with MS according to McDonald’s criteria (McDonald et al., 2001) and nine patients diagnosed with NMOSD according to International NMO diagnostic panel criteria (Wingerchuk et al., 2015). The control CSF samples obtained from healthy individuals were not available, so we recruited CSF samples from twenty-eight patients diagnosed with PTC according to Dandy criteria (Dandy, 1937). As there is limited information about PTC in terms of inflammation, it serves as a suitable unhealthy control, similar to non-inflammatory diseases. For the MS group, patients with the RRMS subtype, and for the NMOSD group, cases diagnosed with the Optic Neuritis and Transverse Myelitis subtypes were recruited. Patients with primary PTC were recruited for the PTC group. CSF samples were collected via the Lumbar Puncture (LP) method and stored at -80°C until performing ELISA analysis. The cases with other autoimmune diseases did not enroll to avoid cross-reactivity.

### 2.2 Quantification of Protein Levels with ELISA

The assays of all the parameters were performed by using the human ELISA kits (Sunred Biological Technology Co., Ltd in Shanghai, China). Before conducting the assays, all CSF samples were carefully thawed at temperatures between 2-8°C and allowed to reach room temperature for 30 minutes. The tests were executed following the supplier company’s instructions.

### 2.3 Statistical Analysis

Statistical analysis was conducted using R Core Team 2023, version 4.3.1. Frequency and percentage were used for categorical-demographic data of the cases and controls. The Kolmogorov-Smirnov test was used to determine the distributional characteristics of continuous variables. One-way ANOVA was used to compare normally distributed data for multiple groups, the LSD was used for pairwise post-hoc comparison, and the Bonferroni correction was applied. Kruskal-Wallis test was used for non-normally distributed data, the Mann-Whitney U test was used as a post-hoc test, and the Bonferroni correction was applied. In descriptive statistics, mean ± standard deviation values were provided for normally distributed characteristics, while median (minimum-maximum) values were given for non-normally distributed characteristics. To detect the relationship between variables, the Pearson correlation test was performed. The statistical significance level was accepted as p < 0.05.

## 3. Results

### 3.1 Demographic and clinical features of patients

Upon analyzing the gender distribution of participants, 86.1% of the patients in the MS group were female and 13.9% were male. 66.7% of the patients were female and 33.3% were male in the NMOSD group. In the unhealthy control group, the percentage of male and female patients was 17.9% and 82.1%, respectively. All participants were aged between 18-65 years. When comparing the mean age of participants based on gender, no statistically significant difference was observed between the mean age of females and males (p= 0.306). The patients’ ages were divided into three groups 18-25, 26-39, and above 40 years. The number of cases aged 18-25 years was higher in the demyelinating diseases group, compared to the PTC group (p=0.025). Gender did not affect age at disease onset (p=1.000). The levels of IgG in the early and late-onset (≤ 35 years and > 35 years, respectively) cases were similar. In terms of EDSS scores, no significant difference was observed between the MS and NMOSD groups. There was no significant correlation between EDSS scores and CAPN1 and CAPN2 levels in the demyelinating disease groups. In Table 1, the demographic and clinical features of the groups are shown.

**Table 1.**
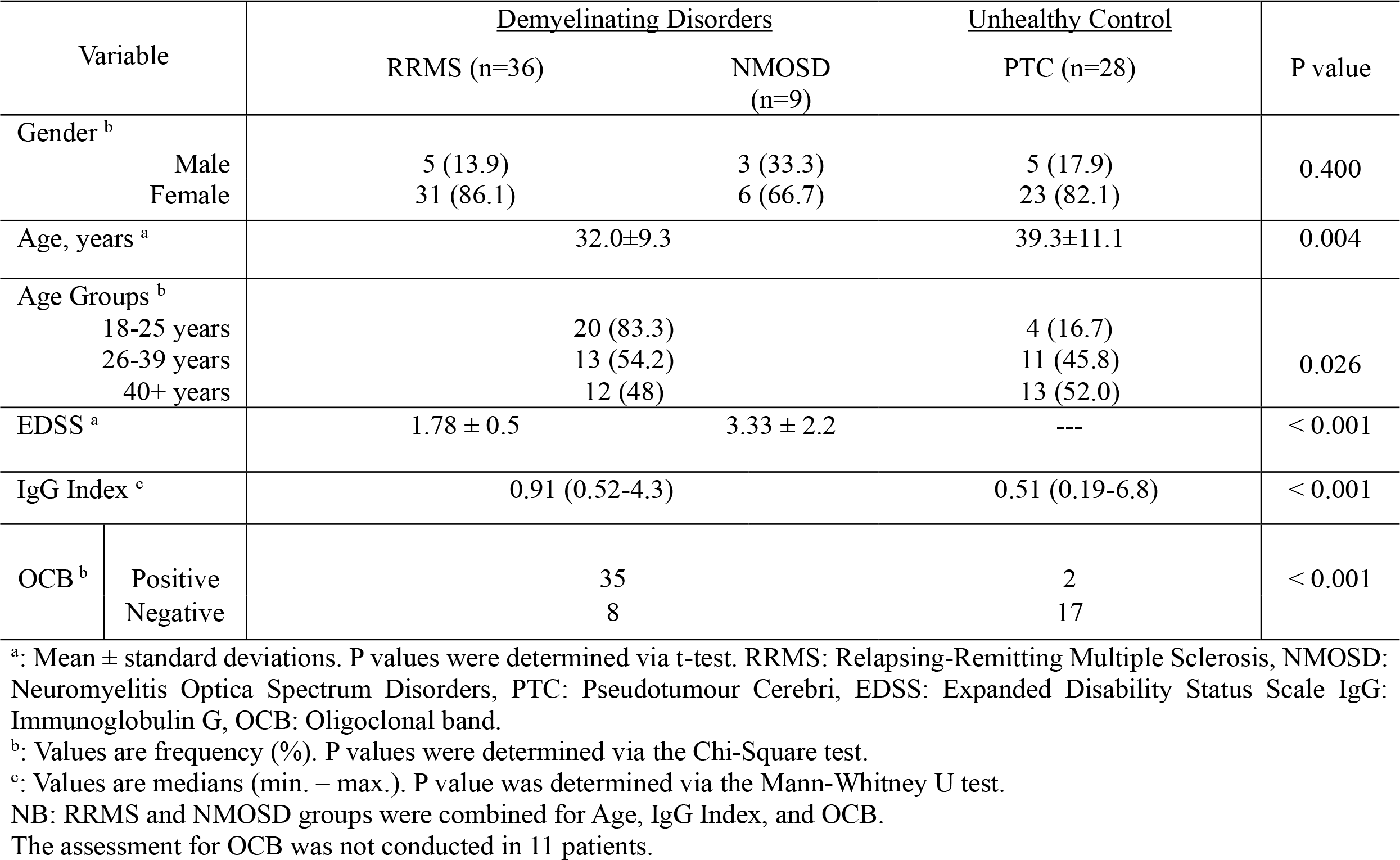
Demographic and clinical characteristics

### 3.2 Assessment of ELISA Results

#### 3.2.1 CAPN1 and CAPN2 levels

Significant differences were found in the levels of CAPN1 in CSF between the MS group and both the PTC group (p=0.001) and the NMOSD group (p=0.002) (Fig. 1). It was a significant increase in the level of CAPN2 in CSF between the MS group and NMOSD group (p=0.002), but no significant difference was found between MS and PTC group. CAPN2 level was higher in the PTC group than NMOSD group (p=0.006).

**Fig. 1.**
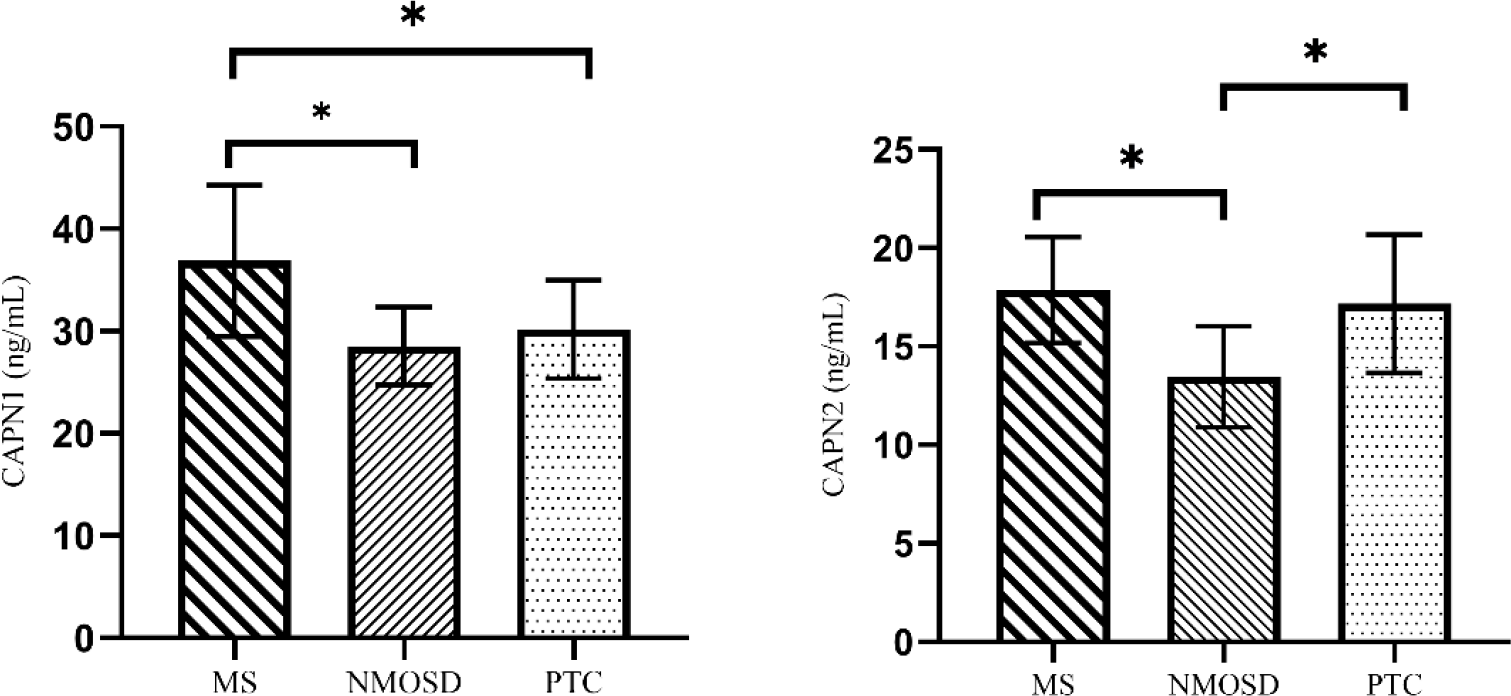
CAPN1 and CAPN2 levels in CSF. The error bar indicates mean ± standard deviation. The group comparisons were performed using one-way ANOVA, and LSD was used as a post hoc test (Bonferroni correction was performed). (*p< 0.001).

**Fig. 2.**
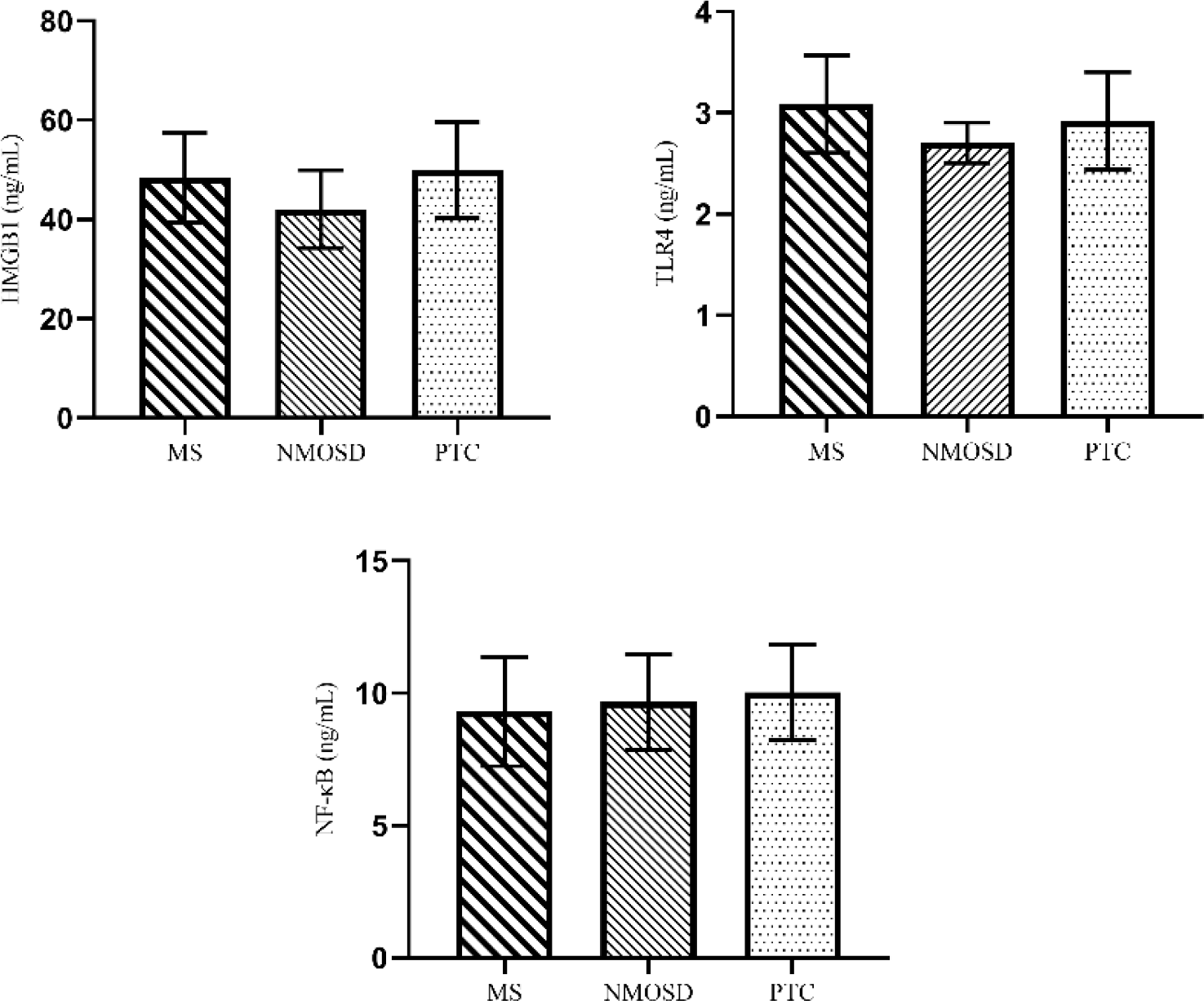
HMGB1, TLR4, and NF-κB levels in CSF. The error bar indicates mean ± standard deviation. The group comparisons were performed using one-way ANOVA, and LSD was used as a post hoc test (Bonferroni correction was performed).

#### 3.2.2 HMGB1, TLR4 and NF-κB levels

No significant differences were found in the levels of HMGB1, TLR4, and NF-κB in CSF between the groups (Fig.2).

#### 3.2.3 IL-17A and IL-37 Levels

No significant difference was found in IL-17A and IL-37 levels among the groups. However, a moderate positive correlation (p=0.001, r=0.472) was found between these two cytokines (Fig. 3.).

**Fig. 3.**
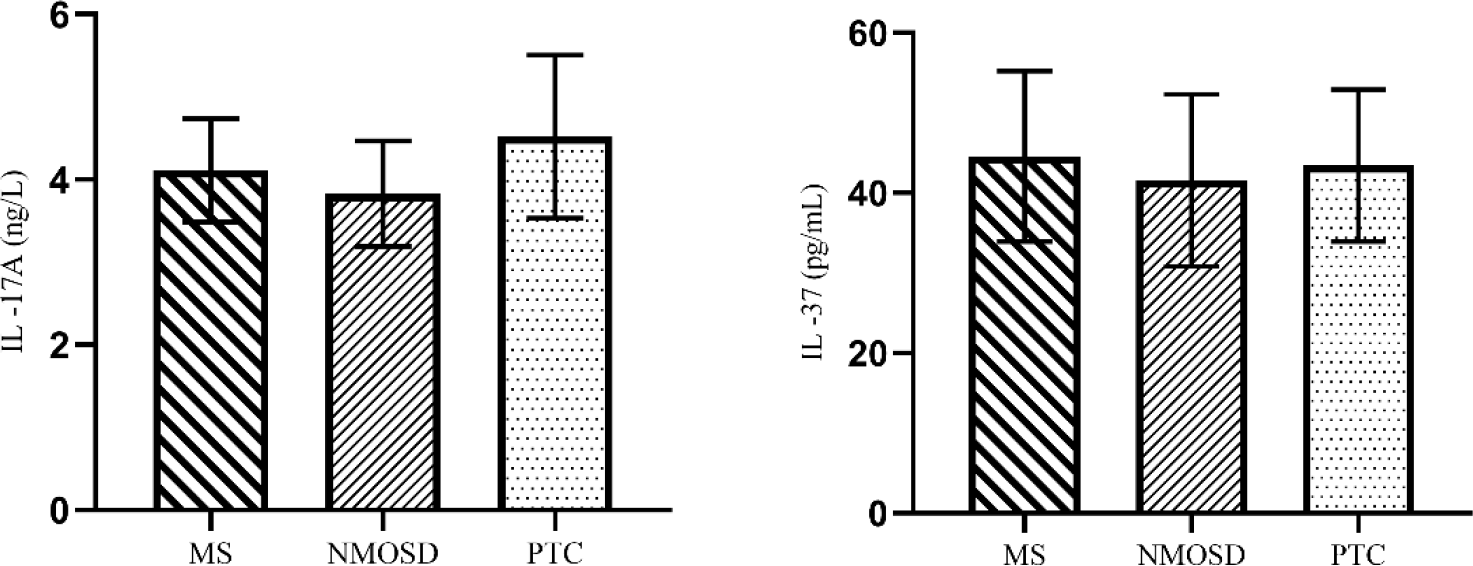
The levels of IL-17A and IL-37 in CSF. The error bar indicates mean ± standard deviation. The group comparisons were performed using one-way ANOVA, and LSD was used as a post hoc test (Bonferroni correction was performed).

#### 3.2.4 Relationship between CAPN1 and CAPN2 with HMGB1/TLR4/NF-κB Signaling Pathway

In the patients with MS, CAPN1 and HMGB1 showed a moderate and positive correlation. Also, TLR4 and NF-κB showed a weak and positive correlation with CAPN1. The correlation matrix, which illustrates the relationship between CAPN1 and HMGB1/TLR4/NF-κB signaling pathway in the MS group, is presented in Figure 4.

**Fig. 4.**
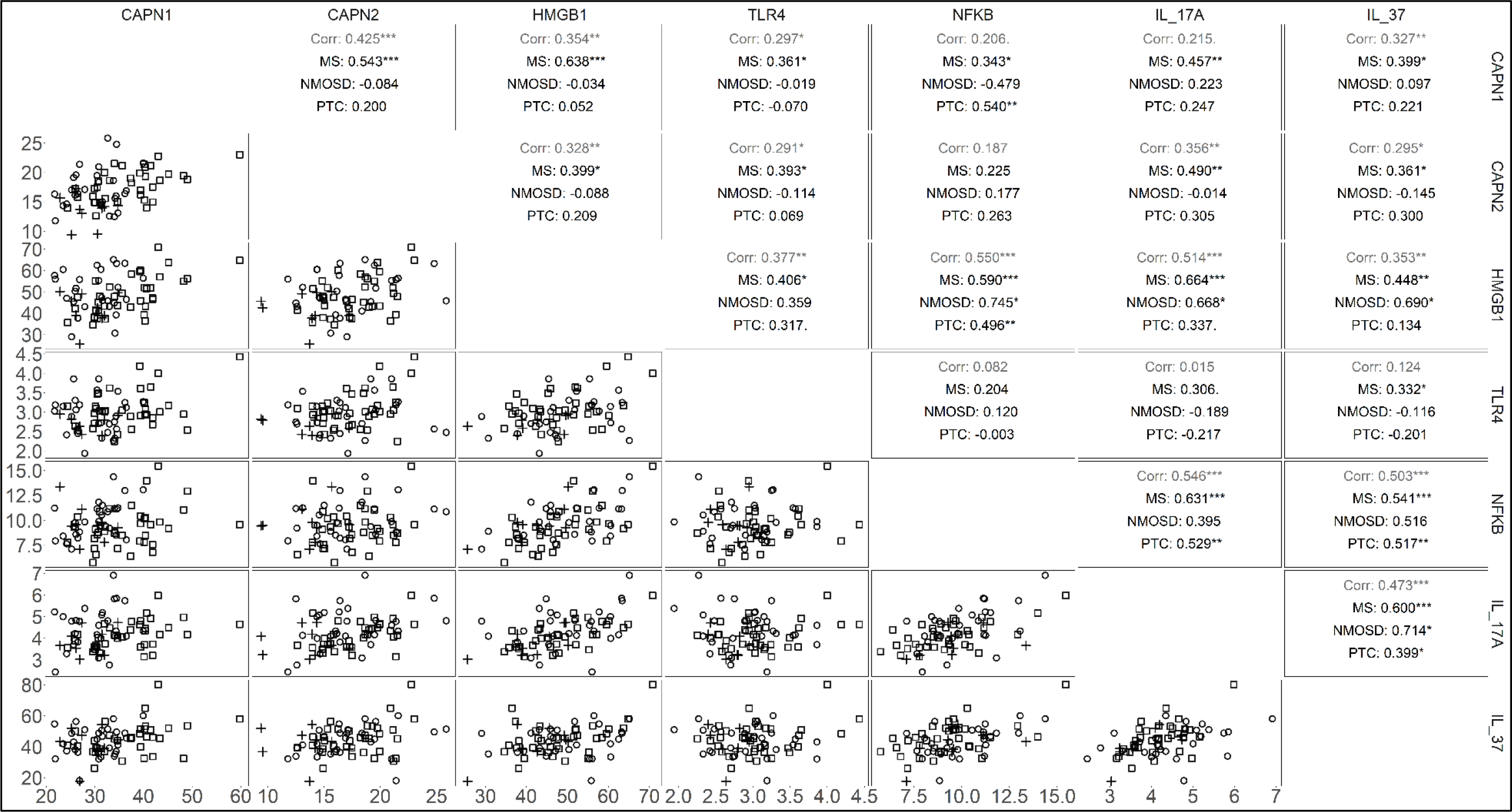
The upper diagonal of the matrix displays the correlation values between CAPN1, CAPN2, and the variables associated with the HMGB1/TLR4/NF-κB signaling pathway, as well as IL-17A and IL-37. The first line represents the overall data, the second line represents the correlation for the MS group, the third line represents the correlation for the NMOSD group, and the fourth line represents the correlation for the PTC group. (*: p < 0.05, **: p < 0.01, ***: p < 0.001). □: MS, +: NMOSD, ○: PTC.)

In patients with MS, a slight positive correlation was noted between CAPN2 and HMGB1, as well as between CAPN2 and TLR4. However, there was no significant correlation found between NF-κB and CAPN2. Figure 4 displays the correlation matrix illustrating the relationship between CAPN2 and the HMGB1/TLR4/NF-κB signaling pathway. Neither CAPN1 nor CAPN2 showed any significant correlation with the HMGB1/TLR4/NF-κB signaling pathway in the NMOSD group, only CAPN1 showed a significant correlation with NF-κB in the PTC group (Fig.4).

#### 3.2.5 Relationship between CAPN1 and CAPN2 with IL-17A and IL-37

Although no correlation was found between CAPN1 and IL-17A, a positive correlation was discovered between IL-37 and CAPN1 (r=0.327, p=0.005). Additionally, positive correlation was observed between CAPN2 and IL-17A and IL-37 (r=0.356, p=0.002, and r=0.295, p=0.011 respectively). In the MS group, there was a moderate positive correlation between CAPN1 and IL17A (r=0.457, p=0.005) and a weak positive correlation between CAPN1 and IL-37 (r=0.399, p=0.016). Similarly, a moderate positive correlation was observed between CAPN2 and IL-17A (r=0.490, p=0.002) and between CAPN2 and IL-37 (r=0.361, p=0.030) in the MS group. However, no significant correlation was found between IL-17A and IL-37 with CAPN1 and CAPN2 in the NMOSD and PTC groups. The graphical results are given in Figure 4.

## 4. Discussion

This pilot study is the first study to compare the levels of CAPN1 and CAPN2 in demyelinating diseases, and their relationship with the HMGB1/TLR4/NF-KB signaling pathway. Upon analysis, it was discovered that patients with MS, NMOSD, and PTC had higher CAPN1 levels than CAPN2. CAPN1 activation is usually associated with neuroprotection, while CAPN2 activation is tied to neurodegeneration (Baudry, 2019). Previous clinical and experimental research has focused mainly on CAPN2 activity in the pathogenesis of MS and NMOSD. CAPN1 is present in the cytoplasm of macrophages, microglia, and astrocytes. Although CAPN2 is primarily found within these cells, it also exists in certain neuronal cytoplasm regions outside of plaques. A recent study examining CAPN1 and CAPN2 expression levels in acute MS lesions found a noteworthy boost in CAPN1 expression relating to axon damage, along with a similar increase in CAPN2. While CAPN1 is present in significantly greater quantities than CAPN2 within acute plaques, both proteins see increases in conjunction with axon damage in these areas. Moreover, research has shown that both calpains exhibit similar levels of expression, even in regions where axon damage is minimal (Baudry et al., 2013; Baudry & Bi, 2016; Gan-Or & Rouleau, 2016; Wang et al., 2020). While previous studies have reported the protective effects of CAPN1, an experimental study found that CAPN1 activation is elevated during the early stages of ALS disease. This increase may be linked to intracellular molecular mechanisms that seek to safeguard neurons from cell death and irreversible damage. According to researchers (Stifanese et al., 2014), an increase in CAPN1 may prevent certain cellular events from occurring, which could otherwise have negative effects. Our study found higher levels of CAPN1 than CAPN2 in newly diagnosed patients with MS during the early stages of the disease, possibly indicating a similar protective mechanism to what was observed in the ALS study. In MS, there is typically a preclinical period of two years, during which neuroinflammation is observed before diagnosis. During this period, both pro-inflammatory and anti-inflammatory cytokines increase (Tzartos et al., 2008). The increase in the levels of CSF CAPN1 in newly diagnosed patients during the early stages could be related to the regulation of cytokine activity, thus it might potentially play a protective role in the neuroinflammation process. However, more research is necessary to determine how CAPN1 and CAPN2 levels change in the later stages of the disease and after treatment before making definitive statements. No significant differences in CAPN1 and CAPN2 levels between groups based on gender or age. However, previous research has shown that CAPN1 levels are higher in female rodents compared to males (Cerghet et al., 2006). The small patient groups in our study may have limited our ability to detect gender differences in calpain levels and gender could be a confounding factor in terms of both CAPN1 and CAPN2 levels or could show an effect with other factors. Besides gender, CAPN1 and CAPN2 levels were not affected by age. However, experimental studies have shown that calpain expression is higher in young rats than in adult rats (Ibrahim et al., 1994). Additionally, another study suggested that there may be a relationship between aging and calpain activity in the brain, indicating that the regional heterogeneity of calpain levels in the brain parallels age-related deterioration (Banay-Schwartz et al., 1994). In MS and NMOSD, advanced age at disease onset may lead to a poor prognosis. Therefore, our findings suggest that further investigation is required to determine the impact of age on calpain levels. More extensive research with larger case and control sample sizes are necessary to establish the correlation between age and calpain levels.

The levels of HMGB1, a marker for inflammation, did not differ significantly among the three groups. Previous research has indicated that NMOSD patients have higher levels of serum HMGB1 compared to MS patients (Wang et al., 2012). A study involving CSF samples reported no significant difference in HMGB1 levels between NMO and MS patients (Wang et al., 2013). Our findings were consistent with this second study. TLR4 and NF-κB levels, both parameters related to HMGB1 activity, were not different between the groups. However, our results did reveal a positive correlation between HMGB1 and TLR4 and NF-κB, which is in line with the mechanism of action of HMGB1.

To evaluate the neuroinflammatory activity of CAPN1, CAPN2, and HMGB1, we analyzed the levels of IL-17A and IL-37, which are known to play a role in the neuroinflammation process. Across all three groups, we observed higher levels of IL-37 compared to IL-17A. While previous studies have suggested that IL-17A may be elevated in the cerebrospinal fluid of MS patients, its detection in serum during different stages of the disease remains controversial. Some studies have explored the role of IL-17A in MS pathogenesis, while others suggest that its detection alone may not be sufficient to determine inflammatory activity (Edwards et al., 2013; Lebrun et al., 2016). We found no correlation between IL-17A and EDSS score and no definite statement about the effect of IL-17A in EDSS. Recent research has shown promise for IL-37 in treating neuroinflammatory diseases, but there is limited data on its efficacy for MS and NMOSD. Previous studies have reported increased IL-37 levels in serum and CSF during acute inflammation (Farrokhi et al., 2015; Li et al., 2013), which is consistent with our findings. These results may suggest that the increased level of anti-inflammatory IL-37 during the preclinical period of MS is reflective of the cytokine/chemokine profile shaping disease activity.

## 5. Conclusions

In this study, CSF samples utilized were obtained from patients during their relapse period. The comparison of the levels of CAPN1 and CAPN2 as well as HMGB1 during remission, was not performed and this is a limitation of our study. Prospective follow-up studies should be conducted on these patients during the remission period. The concept of dichotomy exists concerning the distinct roles of CAPN1 and CAPN2 in neuroinflammation and neurodegeneration, presenting a crucial area of focus for researchers in the development of more precise and effective therapeutic interventions. It is imperative to explore this topic in-depth to gain a better understanding of the mechanistic basis of these diseases and devise targeted treatment strategies to alleviate the symptoms of neuroinflammation and neurodegeneration. It would be advantageous to perform research employing calpain inhibitors, particularly in vitro, to ascertain whether the elevated CAPN1 level we identified in our results is influenced by a neuroprotective mechanism as explained in existing literature, or by a characteristic that contributes to neurodegeneration. Conducting studies using specific inhibitors may produce more conclusive outcomes regarding the direct impact of these calpain species on disease activity and their association with the HMGB1/TLR4/NF-κB signaling pathway.

## Data availability

The data that support the findings of this study are available from the corresponding author upon reasonable request.

## Credit authorship contribution statement

**Firdevs Uluc**: Conceptualization, Methodology, Investigation, Formal analysis, Visualization, Writing – original draft, Writing – review & editing. **Sule Aydin Turkoglu**: Writing – review & editing, Validation, Supervision. **Bihter Gokce Celik**: Conceptualization, Writing – review & editing. **Seyda Karabork**: Writing – review & editing. **Seyit Ali Kayis**: Formal analysis, Validation.

